# A thermodynamic bottleneck in the TCA cycle contributes to acetate overflow in *Staphylococcus aureus*

**DOI:** 10.1101/2024.10.16.618751

**Authors:** Nabia Shahreen, Jongsam Ahn, Adil Alsiyabi, Niaz Bahar Chowdhury, Dhananjay Shinde, Sujata S Chaudhari, Kenneth W Bayles, Vinai C Thomas, Rajib Saha

## Abstract

During aerobic growth, *S. aureus* relies on acetate overflow metabolism, a process where glucose is incompletely oxidized to acetate, for its bioenergetic needs. Acetate is not immediately captured as a carbon source and is excreted as waste by cells. The underlying factors governing acetate overflow in *S. aureus* have not been identified. Here, we show that acetate overflow is favored due to a thermodynamic bottleneck in the TCA cycle, specifically involving the oxidation of succinate to fumarate by succinate dehydrogenase. This bottleneck reduces flux through the TCA cycle, making it more efficient for *S. aureus* to generate ATP via acetate overflow metabolism. Additionally, the protein allocation cost of maintaining ATP flux through the restricted TCA cycle is greater than that of acetate overflow metabolism. Finally, we show that the TCA cycle bottleneck provides *S. aureus* the flexibility to redirect carbon towards maintaining redox balance through lactate overflow when oxygen becomes limiting, albeit at the expense of ATP production through acetate overflow. Overall, our findings suggest that overflow metabolism offers *S. aureus* distinct bioenergetic advantages over a thermodynamically constrained TCA cycle, potentially supporting its commensal-pathogen lifestyle.

## Opinion/ Hypothesis

The gram-positive organism *Staphylococcus aureus* is a frequent colonizer of the human skin and mucosal surfaces of the nose and gut (1). However, it can also invade deeper tissues, causing serious infections such as skin and soft tissue infections, endocarditis and osteomyelitis (2). One of the underlying reasons for its success as a pathogen is its metabolic versatility that allows it to efficiently exploit a variety of niche-specific host nutrients for bioenergetic purposes (3, 4). Yet, when grown in the presence of glucose under conditions of excess oxygen, *S. aureus* appears to execute a seemingly wasteful strategy of excreting substantial amounts of an incompletely oxidized byproduct―acetate, as opposed to fully oxidized CO_2_ (5). This phenomenon is called acetate overflow. Intriguingly, acetate overflow is not unique to *S. aureus but* has been reported to occur in several prokaryotes as well as yeasts that rapidly divide under aerobic growth conditions(6).

At least two major hypotheses have been advanced to explain overflow metabolism. The first involves the proteome allocation hypothesis which argues that energy production through overflow metabolism is more cost-effective than respiration (7–9). The second, membrane real estate hypothesis proposes that overflow metabolism occurs because respiratory capacity is saturated during rapid cell division due to protein crowding on a limited membrane space (10). Here, we show that both the above principles are engaged during rapid growth of *S. aureus* and contribute to acetate overflow. But importantly, their contributions are most accurately reflected only when thermodynamic constraints associated with the TCA cycle of *S. aureus* are accounted for. Although our results indicate that acetate overflow in *S. aureus* is more efficient in generating ATP than the TCA cycle, it is acutely sensitive to the cellular redox status and can easily shift to lactate overflow at the expense of ATP production and growth.

### The cellular redox status impacts acetate overflow in *S. aureus*

Under aerobic conditions, *S. aureus* respires to balance its cellular redox state and generates ATP through oxidative phosphorylation. To assess how cellular respiration impacts acetate overflow, we measured the acetate yield (Y_AC_), defined here as the millimolar concentration of acetate produced per millimolar glucose consumed, in both the wild-type (WT) strain and its isogenic *menD* mutant during aerobic growth. Inactivation of *menD* disrupts menaquinone (MK) biosynthesis, a critical electron carrier in the respiratory chain of *S. aureus*, thereby impairing respiration. Interestingly, analysis of acetate overflow revealed that the acetate yield was significantly higher in the WT strain compared to the *menD* mutant (**Fig 1A-B, Table 1**)(**Supplementary Data 1**). During the exponential phase, the WT strain achieved an acetate yield of approximately 1.48 mM per mM glucose consumed. In contrast, the *menD* mutant had a 74-fold lower Y_AC_ compared to the WT strain. We also observed that anaerobic growth of *S. aureus* under fermentative conditions resulted in diminished acetate overflow, similar to the *menD* mutant (**Fig 1C, Table 1**). These results indicate that active aerobic respiration is essential for acetate overflow during exponential growth.

**Table 1:** Growth parameters of *S. aureus*.

| Growth Condition | Strain | Acetate yield ($Y_{AC}$)<br>(Ac<sub>mM</sub>/ Glu<sub>mM</sub>) | Growth rate<br>($\mu\cdot h^{-1}$) |
| --- | --- | --- | --- |
| (-) O<sub>2</sub> | WT | 0.21 ± 0.06 | 0.46 ± 0.00 |
| (+) O<sub>2</sub> | WT | 1.48 ± 0.10 | 0.56 ± 0.00 |
| | *menD* | 0.02 ± 0.03 | 0.44 ± 0.01 |
| | *atpA* | 1.73 ± 0.04 | 0.58 ± 0.01 |
| | *ccpA* | 1.41 ± 0.01 | 0.57 ± 0.01 |

**Fig. 1:**
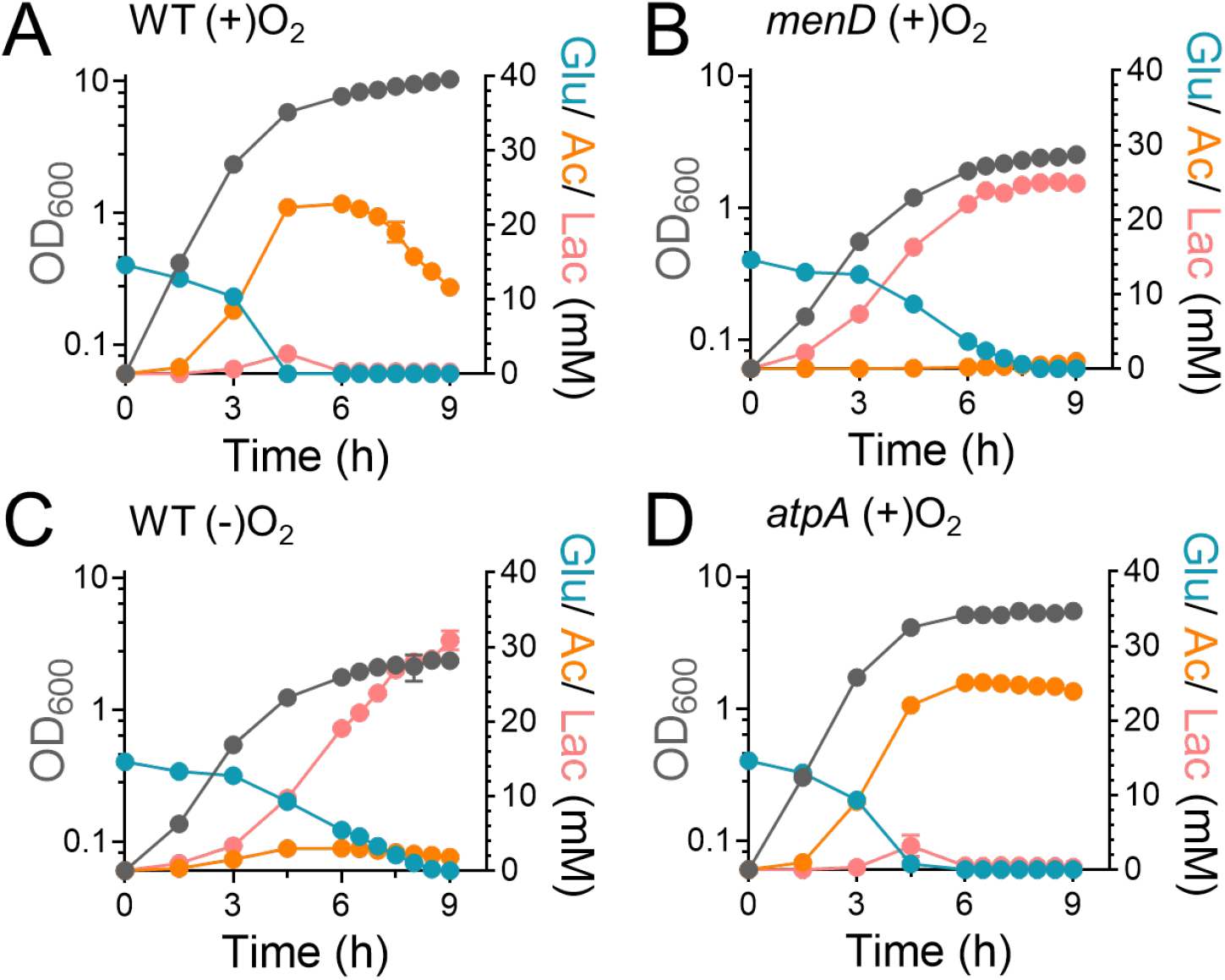
The cellular redox status affects acetate overflow. Glucose, acetate and lactate were assayed in the **(A)** WT, **(B)** *menD*, **(C)** WT (anaerobic) and **(D)** *atpA* mutants during growth (OD_600_). Anaerobic growth of WT strain was carried out under fermentative conditions in the absence of any added nitrate (alternate electron acceptor). n=3, mean ± SD.

When respiration is impaired in the *menD* mutant or when *S. aureus* grows under fermentative conditions, the NAD+/NADH ratio in cells decreases significantly due to reduced transfer of electrons to oxygen via the electron transport chain (ETC). Consequently, pyruvate is used as an alternate electron sink, resulting in its reduction to lactate (**Fig. 1B and 1C**). Additionally, the lack of a functional ETC also compromises ATP production through oxidative phosphorylation as these processes are coupled. To determine the relative importance of redox (NAD+/NADH) maintenance versus ATP production in acetate overflow, we examined the *atpA* (ATPase subunit) mutant, in which ATPase-dependent oxidative phosphorylation is defective but respiration remains functional. Remarkably, the acetate yield of the *atpA* mutant was modestly higher than the WT strain (**Fig. 1D, Table 1**). Thus, the redox balance maintained by a functional ETC is crucial for acetate overflow, whereas ATP generation through oxidative phosphorylation is less critical.

The inactivation of *atpA* had little impact on the aerobic growth rate of *S. aureus* compared to the WT strain (**Table 1**) suggesting that cells rely on alternative pathways for their bioenergetic needs to support growth. Since acetate overflow through the Pta-AckA pathway in *S. aureus* is coupled to ATP production, this suggested that acetate overflow could provide cells with sufficient ATP to maintain its growth rate. Indeed, we found lower growth rates when acetate overflow was not effectively engaged, such as when the WT strain was grown under anaerobic conditions or in the *menD* mutant (**Table 1**) where carbon was redirected to lactate instead of acetate. Overall, these findings suggest that acetate overflow provides an important route to meet the bioenergetic needs of the cell during exponential growth when the cellular redox status is maintained.

### Computational analysis reveals a thermodynamic bottleneck in the TCA cycle that promotes acetate overflow

The use of acetate overflow for energy generation is surprising, given that the predicted ATP gain from 1 mol of acetyl CoA catabolism to acetate is ∼10-fold lower than its complete oxidation to CO_2_ via the TCA cycle. However, in *S. aureus* the TCA cycle is repressed in the presence of glucose due to carbon catabolite repression (CCR) by *ccpA* (11), which may explain why glycolytic flux is primarily directed towards acetate overflow. To test this hypothesis, we determined the acetate yield following *ccpA* inactivation. We reasoned that increased activation of the TCA cycle following *ccpA* inactivation should decrease acetate overflow. Surprisingly, although inactivation of *ccpA* decreased the glucose consumption rate in agreement with previous observations (**Fig. 2A**), the acetate yield of the *ccpA* mutant had only decreased by ∼4.7% compared to the WT strain (**Table 1**). These results suggest that in addition to CCR there may exist other bottlenecks in the TCA cycle of *S. aureus* that contribute to the redirection of carbon flux towards acetate generation.

**Fig. 2:**
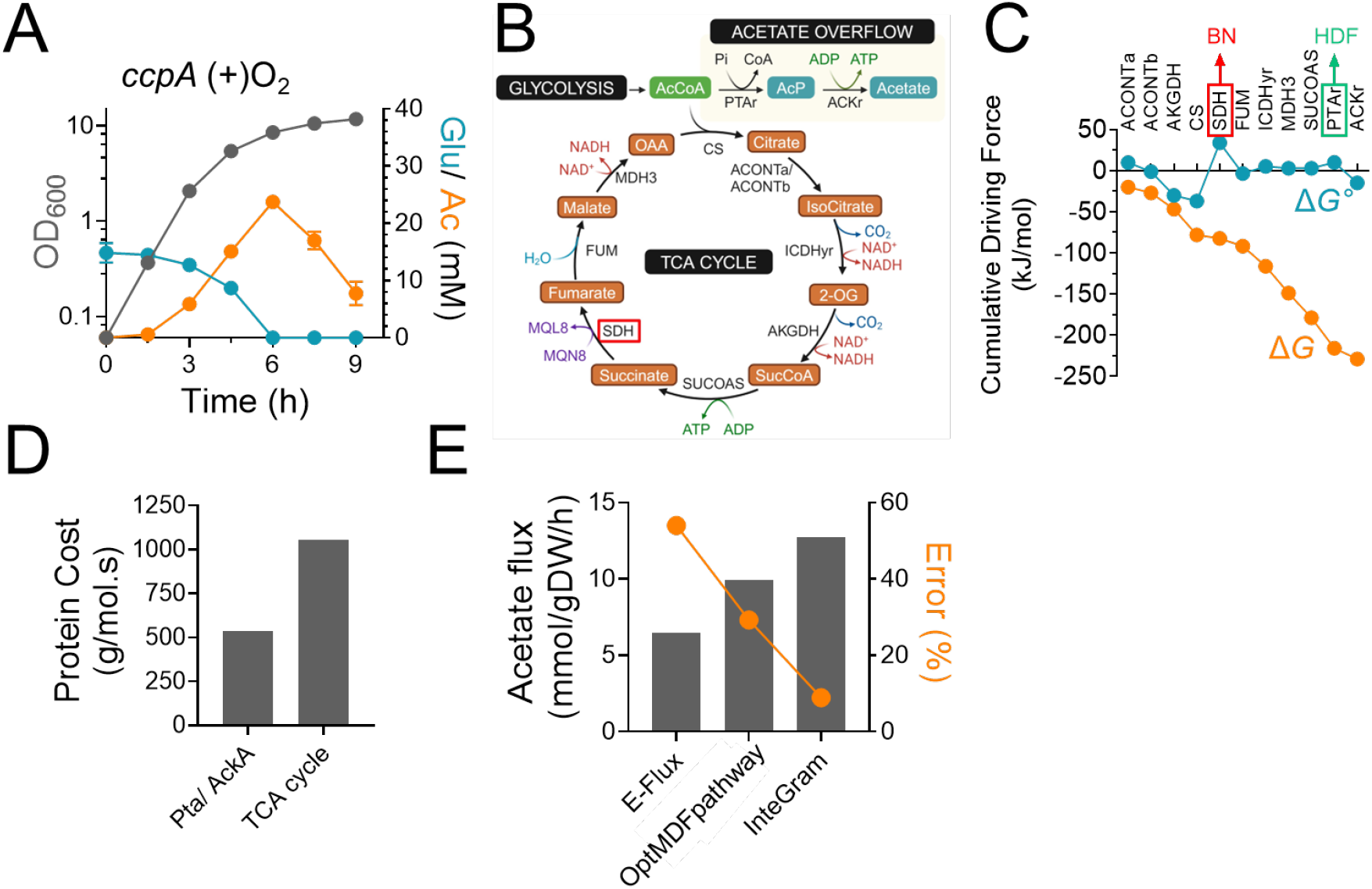
The SDH-catalyzed reaction in the TCA cycle constitutes a bottleneck that leads to acetate overflow. **(A)** Glucose and acetate were assayed in the culture supernatants of the *ccpA* mutant during growth (OD_600_). n=3, mean ± SD. **(B)** Schematic of acetate overflow pathway and TCA cycle. Red boxed enzyme catalyzes bottleneck reaction. **(C)** MDF analysis. Red-boxed enzyme catalyzes the thermodynamic bottleneck (BN) and green boxed enzyme has the highest driving force (HDF). ΔG°, Standard Gibbs free energy; ΔG, Gibbs free energy change was estimated from physiological concentrations of metabolites obtained from experiments and literature survey. **(D)** Protein cost estimation **(E)** Acetate flux estimations from different models were compared against the experimentally observed flux value of 14 mmol/gDW/h. The difference between the estimated and actual acetate flux values were used to determine the % error.

Since metabolic flux through various pathways is governed by thermodynamic constraints, we initially determined the thermodynamic feasibility of the TCA cycle and the acetate overflow pathway (**Fig. 2B**) using the max-min driving force (MDF) framework (12). In the MDF analysis (**Fig. 2C**), the Gibbs free energy change (Δ*G*) was estimated from physiological concentrations of metabolites derived from both experimental data and literature (12). MDF analysis revealed that the acetate overflow pathway (Pta-AckA) had a significantly higher driving force than the TCA cycle, with the phosphotransacetylase (PTAr) reaction showing the highest driving force among the reactions in this pathway (**Fig. 2B-C**). In contrast, a significant thermodynamic bottleneck was observed in the SDH (Succinate dehydrogenase) reaction catalyzed by the *sdhA* gene, which limited the overall driving force of the TCA cycle (**Fig. 2B-C**), making it less favorable. This bottleneck suggests a broader thermodynamic inefficiency within the TCA cycle, potentially driving a shift towards alternative ATP-generating pathways like acetate overflow. Using component contribution analysis, a thermodynamic framework that incorporates both reactant and chemical group contributions to estimate free energy changes, we calculated a minimum standard Gibbs free energy change of +4.9 kJ/mol for the MK-dependent SDH reaction. This contrasts with the -31.1 kJ/mol associated with the ubiquinone-dependent reaction in *E. coli* (13).

Thermodynamic driving force is directly linked to enzyme abundance, with reactions exhibiting lower driving forces requiring a greater enzyme investment to maintain metabolic flux(14). In the presence of a thermodynamic bottleneck, this relationship can significantly increase the enzyme cost, defined here as the total protein mass needed to sustain flux through a pathway. Accordingly, we quantified the enzyme cost for both the TCA cycle and acetate excretion pathway (15). Our calculations revealed that, under aerobic conditions, the TCA cycle incurs a 49% higher protein cost compared to acetate overflow (**Fig. 2D**) (**Supplementary Data 2**). This finding strongly suggests that acetate overflow is preferred due to its lower metabolic burden on the cell.

### Membrane crowding contributes to acetate overflow

In addition to these constraints, physical limitations within cells also contribute to acetate overflow. The membrane real-estate hypothesis suggests that during rapid aerobic growth, the available membrane space becomes saturated with proteins, including those involved in nutrient uptake. This crowding may restrict the cell’s ability to carry out oxidative phosphorylation, further driving the shift toward acetate overflow as an alternative bioenergetic strategy.

To investigate this hypothesis, we initially assessed the gene expression of WT strain during aerobic exponential growth **(Supplementary Data 3)** and integrated the data into our previously published *S. aureus* genome-scale metabolic model (GEM) (16).Before integration, the model was refined to ensure 100% stoichiometric consistency and mass balance. We then applied several algorithms, including iMAT (Integrative Metabolic Analysis Tool) (17), RIPTiDe (Reaction Inclusion by Parsimony and Transcript Distribution) (18), EXTREAM (Expression distributed REAction flux Measurement) (19), and E-Flux (20) to incorporate the transcriptomic data into the GEM. These methods helped contextualize the model by refining reaction constraints based on gene expression data, thereby improving its accuracy in predicting metabolic behavior. Among these methods, E-Flux produced the most accurate predictions, with acetate fluxes closer to experimental values compared to the other algorithms. However, despite its improved accuracy, the error percentage between the predicted and experimental acetate fluxes remained high at 54%. To improve the predictability of this contextualized model, we subsequently implemented OptMDFpathway algorithm (21) with the E-flux model. OptMDFpathway incorporates thermodynamic constraints into pathway analysis, optimizing the distribution of metabolic fluxes to maximize the driving force of reactions while minimizing thermodynamic bottlenecks (**Supplementary Data 4**). Implementation of OptMDFpathway further reduced the error percentage to 29% and closed the gap between the predicted and experimentally observed rates of acetate flux.

Although OptMDFpathway captured acetate overflow well, we still lacked a framework to account for the effect of the surface area of membrane-bound enzymes to test the membrane real-estate hypothesis. To address this, we developed a new algorithm, InteGraM. The algorithm calculated the surface area of membrane-bound enzymes using molecular weight-based empirical equations and constrained the surface area-weighted sum of fluxes of the membrane-bound reactions in the OptMDFpathway formulation. This additional constraint increased acetate flux and reduced the error to ∼9% when the sum of fluxes for membrane-bound reactions was constrained (**Fig. 2E**). Taken together, these results strongly suggest that membrane crowding may further constrain an already flux-limited TCA cycle, reducing ATP production and leading to acetate overflow as a preferred mechanism for energy generation in *S. aureus*.

### Succinate to fumarate reaction constitutes a thermodynamic bottleneck in the TCA cycle

Given that the computational analysis indicated reduced TCA cycle flux from succinate to fumarate is critical for acetate overflow, we investigated whether the *sdhA* catalyzed a bottleneck reaction in the TCA cycle. We utilized liquid chromatography tandem mass spectrometry (LC-MS/MS) to determine the mass isotopologues distribution (MID) of succinate and fumarate derived from^13^C_6_-glucose in *S. aureus* (**Supplementary Fig. 1, Supplementary Data 5**). We specifically focused on two isotopologues (M+2 and M+4) of succinate and fumarate to assess flux through the SDH reaction. When grown in media supplemented with ^13^C_6_-glucose, *S. aureus* generates ^13^C_2_-acetylCoA which can enter the TCA cycle and react with either unlabeled oxaloacetate to generate ^13^C_2_-citrate or with ^13^C_3_-oxaloacetate to form ^13^C_5_-citrate (**Fig. 3A**). Due to subsequent decarboxylation reactions in the TCA cycle, ^13^C_5_-citrate is converted to ^13^C_4_-succinate (M+4) and then to ^13^C_4_-fumarate (M+4) whereas ^13^C_2_-citrate retains both ^13^C-labeled carbon atoms (M+2) when it is eventually converted to fumarate (**Fig. 3A**).

**Fig. 3:**
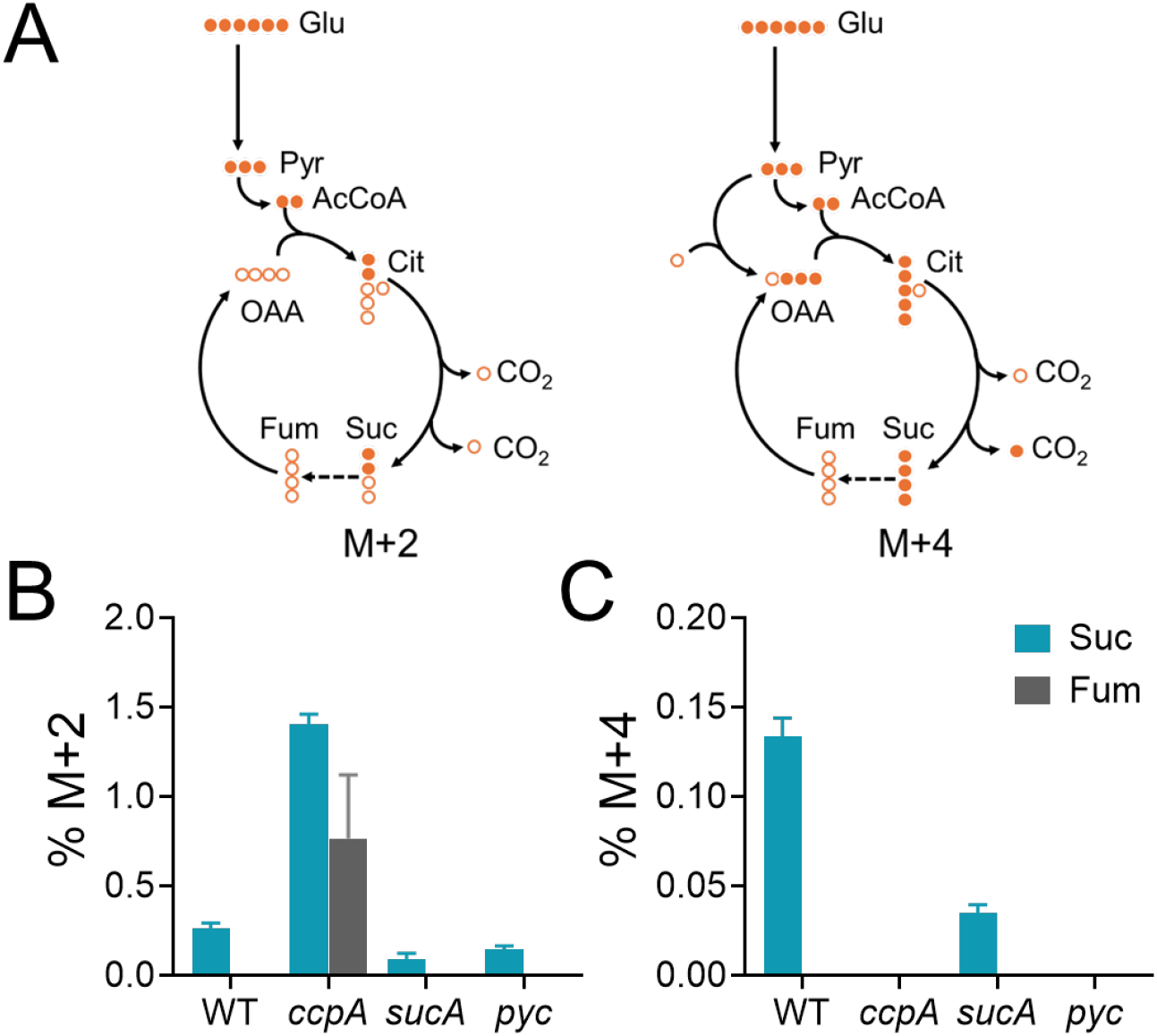
SdhA catalyzes a bottleneck reaction in the TCA cycle. **(A)** Carbon transition scheme resulting in M+2 and M+4 isotopologues. Dashed arrow indicates a bottleneck reaction. **(B)** % M+2 **(C)** % M+4 isotopologues of succinate (blue bars) and fumarate (grey bars) were determined by LC-MS/MS following growth of *S. aureus* in TSB media supplemented with U-^13^C-glucose (n=3, mean ± SD).

We used isogenic *sucA* and *pyc* mutants as controls in these ^13^C-tracing experiments to validate the identity of the M+2 and M+4 isotopologues of both succinate and fumarate. As expected, the inactivation of *sucA* partially reduced the flux from 2-Oxoglutarate to succinate in the WT strain, indicated by reduced levels of the M+2 and M+4 succinate isotopologues (**Fig. 3B**). In contrast, the *pyc* mutation completely eliminated M+4 isotopologues of succinate and fumarate due to the absence of 2-oxaloacetate production from phosphoenolpyruvate (PEP) (**Fig. 3C**).

Upon validation, we quantified the abundance of M+2 and M+4 isotopologues of succinate and fumarate relative to their total pool in cells. LC-MS/MS analysis revealed that the M+2 and M+4 succinate constituted just 0.2% and 0.13% of the total intracellular succinate pool of the WT strain, respectively (**Fig. 3B-C**), suggesting that the flux through the citrate node of the TCA cycle was limited during rapid exponential growth in media containing glucose. Importantly, we did not detect either M+2 or M+4-labeled fumarate from their corresponding succinate isotopologues in the WT strain (**Fig. 3B-C**), which is consistent with a bottleneck in the *sdhA*-catalyzed reaction. This bottleneck was partly relieved in the *ccpA* mutant where M+2 succinate and fumarate increased to 1.4% and 0.76% respectively (**Fig. 3B**). We did not observe a corresponding increase in M+4 succinate (**Fig. 3C**), presumably because *ccpA* positively regulates *pyc* expression (22). Consistent with this hypothesis, the M+4 succinate pool was also depleted in the *pyc* mutant (**Fig. 3C**). Collectively, these results suggest that the conversion of succinate to fumarate constitutes a bottleneck in the TCA cycle which could lead to acetate overflow.

## Conclusions

Overflow metabolism typically refers to the incomplete oxidation of glucose to acetate or lactate by cells even when oxygen is abundantly present. It is a seemingly inefficient way to utilize carbon for bioenergetic purposes.

We propose that a thermodynamic bottleneck in the conversion of succinate to fumarate in the TCA cycle drives acetate overflow in *S. aureus*. Isotope tracing studies using U-^13^C_6_-glucose revealed reduced flux through the TCA cycle during rapid cell division and supported a bottleneck in the conversion of succinate to fumarate, as neither M+2 nor M+4 isotopologues were detected for fumarate from their corresponding succinate species. The enzyme SDH catalyzes this reaction, which is coupled to MK reduction in the respiratory chain. The bottleneck arises from the low midpoint potential of the MK redox couple (∼ -72 mV) (23), requiring a substantial amount of both succinate and SDH to drive the reaction efficiently towards fumarate. In contrast, gram-negative organisms like *E. coli* that utilize ubiquinone (UQ) as their primary aerobic respiratory quinone can easily drive the same reaction following interaction with SDH due to the substantially positive redox potential of UQ (mid-point potential of UQ/UQH2 ∼110 mV) (24). When *E. coli* is forced to respire using MK due to a mutation that prevents UQ biosynthesis, these cells are unable to efficiently grow aerobically and disproportionately rely on acetate overflow to meet their bioenergetic needs (25). These findings strongly support the hypothesis that the thermodynamic limits of the *sdhA* catalyzed reaction in the TCA cycle, drives acetate overflow in *S. aureus*.

Given the thermodynamic constraint in TCA cycle, it becomes imperative to ask if acetate overflow through the Pta-AckA pathway still constitutes a wasteful strategy for ATP generation. Even though the ATP yield from 1 mol of glucose catabolized through the TCA cycle far exceeds that of acetate overflow, our analysis shows that the thermodynamic constraint on the SDH reaction considerably decreases flux through this node of the TCA cycle, making it more economical for *S. aureus* to support ATP generation through acetate overflow pathways than the bacterial ATP synthase. Our calculations show that ATP flux through acetate overflow is ∼3.5-fold more than TCA-dependent respiration (ATP flux of *12mmologDW*^−1^*h*^−1^ vs 3.5 *mmologDW*^−1^*h*^−1^, respectively). From a separate perspective, when protein costs are considered after factoring in the thermodynamic constraint in the TCA cycle, the cost for acetate overflow pathway is 536 *g/(mols*^−1)^, whereas respiration supported through TCA cycle activity had a much higher enzyme cost of 1055 *g/(mols*^−1)^. This suggests that the TCA/respiration route would require about 2-fold more investment of enzymes per unit of ATP generated compared to acetate overflow. Overall, these findings suggest that *S. aureus* may favor acetate overflow since it is more advantageous and cost effective for ATP production than the TCA cycle.

It is intriguing to consider why CcpA represses the TCA cycle if a thermodynamic bottleneck already limits its flux. Indeed, isotope tracing revealed only a modest increase in TCA cycle flux following *ccpA* inactivation which corresponded to a marginal decrease in acetate yield. We propose that the CcpA-dependent carbon catabolite repression of the TCA cycle may primarily help limit the production of energetically costly enzymes, which would otherwise be wastefully produced for a flux-restricted pathway. Thus, the impact of CcpA on TCA cycle activity may be viewed more as a fine-tuning function rather than an on/off switch.

Finally, it is surprising that *S. aureus* has evolved to only utilize MK for respiration when clear advantages for UQ exist as far as economical carbon utilization. Our results show that the use of MK underlies overflow metabolism and provides *S. aureus* with the ability to redirect carbon towards ATP production or redox regulation in a rapid efficient manner. This may ultimately aid its ability to switch from an oxygen rich environment on the skin to more hypoxic environments deep within tissues, thus allowing a smooth transition between its commensal/pathogen lifestyle in the human host. It is tempting to speculate that organisms that utilize UQ may have a relatively limited spare capacity for carbon redirection towards redox control as flux through TCA cycle may compete out those of redox pathways. Thus, our findings suggest that the use of acetate overflow over the TCA cycle not only provides an alternate economical source of ATP for *S. aureus*, but also likely contributes to its metabolic versatility.

## Data availability

The materials and method can be found in the Supplementary file. All codes have been deposited in GitHub (https://github.com/ssbio/Staph).

## Acknowledgments

We gratefully acknowledge funding support from the National Institute of Health (NIH) R35 MIRA grant (5R35GM143009) to RS, 2P01A1083211 to KB and NIH/NIAID R01AI125588 and 2P01A1083211 (Metabolomics Core) to VCT, and Nebraska Collaboration Initiative (NCI) Grant (21-1106-6009) to KB, RS, and VT. Mass spectrometry analyses were performed by the University of Nebraska Medical Center Multiomics Mass Spectrometry Core Facility (RRID: SCR_012539).

